# Parsimonious cell co-localization scoring for spatial transcriptomics

**DOI:** 10.64898/2026.02.09.704806

**Authors:** Ian K. Gingerich, Hildreth Robert Frost

**Affiliations:** Department of Biomedical Data Science, Geisel School of Medicine, Dartmouth College, Hanover, NH, USA

## Abstract

Spatial transcriptomics (ST) preserves tissue architecture while profiling gene expression, motivating methods that quantify whether annotated labels (such as cell types) preferentially co-occur in local neighborhoods. We introduce the Neighborhood Product Co-localization (NPC) score, a simple per-cell metric computed on a pruned spatial neighbor graph: for a set of *m* ≥ 2 labels, NPC is the product of their neighborhood proportions, optionally normalized by expected co-occurrence under independence and paired with permutation-based significance testing. NPC is interpretable (maximized under balanced neighborhoods), efficient to compute, and extends naturally from pairwise to multivariate microenvironment definitions. Using a mouse ovary MERFISH dataset, we show that NPC complements established Squidpy co-occurrence and neighborhood enrichment analyses by localizing co-localization hotspots in tissue space, recapitulating prominent global associations, and highlighting spatially restricted niches such as follicle boundaries; we further demonstrate multivariate NPC scoring by identifying coordinated endothelial–stroma–theca co-localization. Overall, NPC provides a practical framework for interpretable, single-cell resolution co-localization analysis in ST cohorts.

## 1 Introduction

Spatial transcriptomics (ST) technologies measure gene expression while preserving the two-dimensional organization of tissue, enabling joint analysis of molecular state and spatial context at cellular resolution[1, 2]. By integrating ST data with cell-type annotation, one can ask not only which cell types are present, but also how they are arranged and intermixed across microenvironments such as immune infiltrates, stromal niches, vasculature, and epithelial compartments [3]. Cell-type co-localization analyses address these questions by quantifying whether particular cell types preferentially occur near one another (or avoid each other) beyond what would be expected by chance, thereby providing interpretable proxies for putative cell–cell interactions, tissue remodeling processes, and disease-associated spatial architectures.

A wide range of approaches has been proposed to quantify spatial association, spanning simple descriptive statistics through increasingly model-rich or computationally intensive frameworks. At the simplest end are graph-based summaries such as Squidpy’s neighborhood enrichment and co-occurrence probability ratio, which quantify whether pairs of cell types are observed as neighbors (or within increasing radii) more often than expected under random label permutations [4]. Classical spatial statistics—including Moran’s I for spatial autocorrelation and Ripley’s K/L-family functions for attraction/repulsion in marked point patterns—similarly provide principled, scale-aware summaries of spatial structure [5][6][7]. More elaborate methods build empirical null models across multiple length scales (e.g., CRAWDAD’s tile-based label shuffling with binomial proportion testing) and summarize scale-dependent association as “trend” curves [8]. Others explicitly treat images as marked point processes and reduce differences between observed and Poisson-expected L-curves to a single per-image co-localization score that can be tested across experimental conditions using linear or mixed-effects models (e.g., `spicyR`; related functional/point-pattern tools include `spatstat`, `spatialFDA`, and lattice-based methods in `Voyager`/`spdep`) [9–12]. Finally, deep learning and spatial mapping approaches can define co-localization via shared mapping probabilities of single cells to spatial locations (e.g., constructing a cell–cell co-localization matrix from normalized spatial assignment distributions as in DeepCOLOR), with significance assessed via permutations [13].

While these approaches are powerful and often provide statistical guarantees or multi-scale interpretability, they can be less convenient for large ST cohorts when the goal is to (i) obtain an interpretable, local, per-cell readout, (ii) screen many cell-type combinations, (iii) score more than two cells at once, and (iiii) maintain computational tractability. In practice, scale-sweeping methods (e.g., grid/tiling permutations or full distance-based summaries) can be expensive and yield outputs that are primarily pairwise or image-level curves, making it harder to localize where in tissue an association occurs or whether it is driven by a small subset of regions/samples; point-process approaches typically require careful specification of observation windows and edge correction; and mapping-based or deep-learning-derived co-localization can scale quadratically with cell number and may depend on additional modeling assumptions and tuning choices. In contrast, our Neighborhood product co-localization (NPC) score is a model-light function of neighborhood compositions on a standard spatial neighbor graph: it produces per-cell values with a clear maximum under balanced neighborhoods, supports straightforward abundance normalization and permutation testing, and extends naturally from pairwise to multivariate microenvironment definitions (e.g., endothelial+CD8+myeloid), enabling efficient, interpretable “hotspot” mapping and sample-level summarization across conditions.

## 2 Methods

### 2.1 Neighborhood Product Co-localization (NPC) score

#### 2.1.1 Spatial neighbor graph construction and pruning

To quantify spatial co-localization between sets of annotated cell types (any label would work), our approach computes a per-cell co-localization score for a specified set of cell labels. For each sample, a spatial *k*-nearest neighbor graph is constructed from cell centroid coordinates (e.g., `squidpy.gr.spatial_neighbors` with `delaunay = False` and `n_neighs = 50`). Long-range edges are removed using a link-pruning procedure (e.g., `CellCharter remove_long_links`)[14] and self-edges are excluded. Cells with zero remaining neighbors after pruning are excluded from score computation and assigned missing values for the co-localization scores. Let *n* be the number of retained cells after pruning. Let *A* ∈ {0, 1}^*n×n*^ denote the pruned adjacency matrix, where *A*_*ij*_ = 1 if cell *j* is considered a spatial neighbor of cell *i* (and *A*_*ij*_ = 0 otherwise). Define the row-normalized weight matrix *W* ∈ ℝ^*n×n*^ by

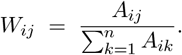

Thus, *W*_*ij*_ represents the proportion of cell *i*’s neighborhood attributed to neighbor *j*.

#### 2.1.2 Binary indicators of cell identity

For each label *c*, define a binary indicator vector *x*^(*c*)^ ∈ {0, 1}^*n*^ by

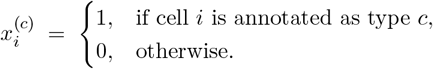

#### 2.1.3 Local neighborhood proportions (“spatial lags”)

For each cell *i* and label *c*, the neighborhood proportion (spatial lag) is defined as

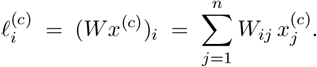

These quantities can be interpreted as the fraction of cell *i*’s neighbors belonging to label *c*.

#### 2.1.4 Multivariate co-localization score (more than two labels)

Let *C* = {*c*_1_,…, *c*_*m*_} be a set of *m* ≥ 2 labels of interest. The multivariate co-localization score for the set *C* and cell *i* is defined as the product of neighborhood proportions:

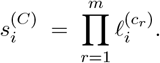

This score is high when cell *i* has a neighborhood that is simultaneously enriched for all cell labels in *C*.

##### Maximum under an even split

By the arithmetic–geometric mean inequality, for nonnegative 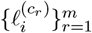 with fixed sum, the product 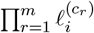 is maximized when all terms are equal. In particular, if we consider neighborhoods in which the mass is concentrated on the *m* labels in *C* (i.e. 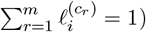), then the maximum occurs at an even split

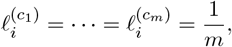

yielding the maximum score

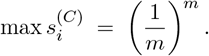

Thus, the score is largest when the local neighborhood is balanced across the specified labels rather than dominated by one.

##### Relationship to entropy-like diversity measures

Let 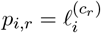 denote the neighborhood composition over the *m* types in *C* (with 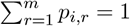 under the restriction above). The quantity

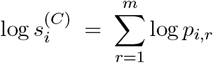

is maximized at the uniform distribution *p*_*i,r*_ = 1*/m*. This behavior is qualitatively similar to entropy-based measures of neighborhood diversity, which are also maximized under an even split. Unlike Shannon entropy, the product score emphasizes the simultaneous presence of *all* specified types; if any *p*_*i,r*_ = 0, then 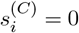.

##### Abundance normalization

To account for the global abundance of each label, we define a normalized score by dividing by the expected product under independence:

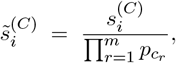

where 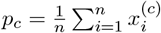 is the marginal frequency of type *c*. Normalization is applied only when 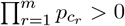.

##### Permutation-based significance testing

Empirical *p*-values are computed using a label-permutation null model. Specifically, cell-type labels are randomly permuted across the *n* retained cells *B* times (e.g., *B* = 1000), preserving the spatial graph and the marginal counts of each label. For permutation *b* ∈ {1,…, *B*}, recompute

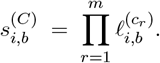

The one-sided empirical *p*-value for each cell is then

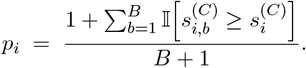

As above, 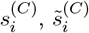, and *p*_*i*_ can be stored for downstream analysis.

#### 2.1.5 Pairwise co-localization as a special case

The pairwise version of the method is obtained by setting *m* = 2 and *C* = {*c*_1_, *c*_2_}, yielding 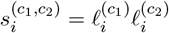, with the corresponding normalization and permutation testing defined analogously.

### 2.2 Comparison techniques for spatial cell-type association

To benchmark and contextualize the proposed Neighborhood Product Co-localization (NPC) score, we additionally computed two established graph-based spatial association summaries implemented in Squidpy [4]. First, we computed the co-occurrence probability ratio using `squidpy.gr.co_occurrence`, which evaluates whether a query label is enriched or depleted within increasing radii around a conditioning label, relative to the query label’s global frequency. Second, we computed global neighborhood enrichment using `squidpy.gr.nhood_enrichment`, which quantifies whether edges between label pairs occur more or less frequently than expected under label permutations on a fixed spatial neighbor graph, and reports an enrichment *z*-statistic for each label pair. These comparison methods provide complementary global and/or scale-dependent summaries, but do not yield per-cell co-localization scores.

## 3 Results

We applied our local co-localization score to the mouse ovary MERFISH spatial transcriptomics dataset (5)[15], which provides dense cellular sampling and well-defined anatomical compartments, making it suitable for evaluating cell-type spatial organization. As visualized in (Fig. 1A), the annotated cell types form clear structural domains and boundaries, motivating complementary global and local spatial statistics to quantify these patterns.

**Figure 1.**
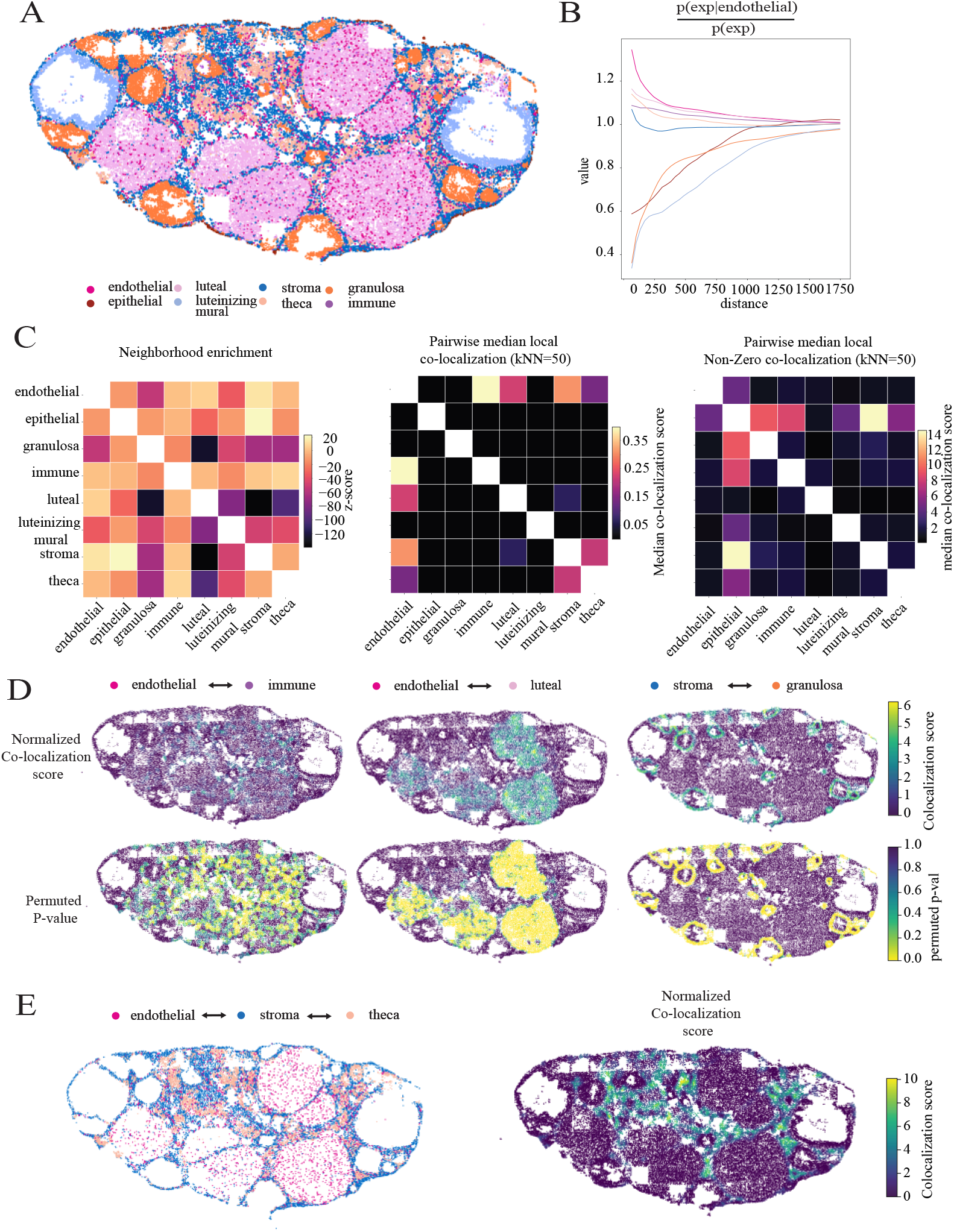
Spatial statistics computed on mouse Ovary ST data (MERFISH). A: Spatial skatterplot colored by cell type. **B:** Co-occurrence of cell types compared to endothelial cells. **C:** Heatmap of global neighborhood enrichment Z-statistic (left), median local co-localization scores (middle), and median non-zero local co-localization scores (right). **D:** Normalized co-localization scores (top row) and permuted p-values (bottom row) for endothelial-immune, enothelial-luteal, and stroma-granulosa co-localization visulized spatially. **E:** Example of endothelial-stroma-theca cell co-localization colored by cell type (left) and normalized co-localization score (right).

Using established Squidpy routines as comparison techniques, we first computed the co-occurrence probability ratio with endothelial cells as the conditioning population. This analysis revealed strong short-range enrichment of endothelial–endothelial proximity as expected, together with elevated co-occurrence of endothelial cells with luteal and theca populations at small radii (Fig. 1B). We next applied Squidpy’s global, graph-based neighborhood enrichment as a second comparison method, which highlighted prominent pairwise associations across the tissue: epithelial–stromal neighborhoods exhibited a strong positive enrichment signal, while granulosa–luteal and stromal– luteal pairings were strongly depleted (Fig. 1C, left).

We then applied our proposed NPC score to compute per-cell local co-localization values for all cell-type pairs and summarized these scores to obtain an aggregated view of local neighborhood composition. The median NPC matrix recapitulated the dominant epithelial–stromal relationship observed by global enrichment, while mphasizing other pairings that were less apparent in the global enrichment heatmap, including epithelial–granulosa and epithelial–immune co-localization (Fig. 1C, middle). When restricting aggregation to non-zero NPC scores, the resulting matrix further prioritized interactions where co-localization occurs in spatially localized tissue niches rather than being diluted by large numbers of cells with zero signal (Fig. 1C, right), providing a complementary view of “where present, how strong” co-localization.

A key advantage of the proposed method is that it yields per-cell scores that can be mapped back into tissue space. Spatial visualization of normalized endothelial–immune, endothelial–luteal, and stroma–granulosa scores revealed distinct anatomical patterns, with high-scoring regions and corresponding permutation-derived significance values aligning with interpretable tissue structures, including follicles and follicle boundaries (Fig. 1D). Finally, we demonstrate the multivariate extension of the score by examining endothelial–stroma–theca co-localization: plotting the three cell types alongside the corresponding normalized multitype score highlights spatially restricted regions where these populations jointly co-occur, consistent with localized microenvironments at compartment interfaces (Fig. 1E). Collectively, these results show that our local co-localization score complements global spatial association measures by preserving interpretability at single-cell resolution while remaining compatible with permutation-based significance assessment and extension to multitype neighborhood definitions.

## 4 Discussion

The co-localization score introduced here is intended as a practical complement to existing spatial association analyses, with particular strengths for cohort-scale studies where interpretation and visualization at single-cell resolution are central. Unlike methods that primarily return cell-type–pair summaries (e.g. enrichment matrices or scale-dependent association curves), the proposed metric produces a per-cell readout. This enables direct mapping of co-localization ‘hotspots’ back onto tissue coordinates, facilitating qualitative inspection alongside histology and quantitative characterization of where signal concentrates (e.g. whether it is diffuse, confined to specific compartments, or driven by a small number of focal regions). Because the score is computed for every cell, it also supports downstream analyses that are naturally phrased in terms of distributions (e.g. the fraction of cells exceeding a threshold, spatial clustering of high-scoring cells, or comparisons of score distributions across conditions), which are cumbersome to derive from global pairwise summaries alone.

A second advantage is that the score directly targets the simultaneous presence of multiple cell types within the same local neighborhood. Many widely used spatial association measures are fundamentally pairwise, quantifying whether two labels tend to occur nearby more often than expected. In contrast, the product-of-proportions formulation extends cleanly to multivariate definitions (e.g. endothelial+CD8 T+myeloid) and naturally emphasizes neighborhoods where all specified types are represented. The score is maximized by an even split among the chosen types, providing an intuitive interpretation related to neighborhood mixing/diversity: high values correspond to balanced multi-lineage microenvironments rather than neighborhoods dominated by one type with occasional contact to another. This multivariate capability is particularly useful when the biological object of interest is not a single interaction pair but a structured niche or domain (e.g. immune–vascular–stromal interfaces) that would otherwise require piecing together multiple pairwise tests.

Third, the metric is deliberately interpretable. The neighborhood proportions have a clear meaning—“what fraction of my neighbors are of type *c*?”—and the co-localization score is a simple combination of these quantities. This transparency makes it easier to reason about how preprocessing decisions (cell-type annotation granularity, neighbor-graph construction, pruning of long edges) affect results. At the same time, the approach retains a natural handle on spatial scale without requiring explicit computation across many radii: scale can be explored by adjusting the same neighbor graph (e.g. *k* in kNN, distance cutoffs, or link-pruning parameters) while keeping the score definition unchanged. In practice, this often provides sufficient control over locality while avoiding the computational overhead and added complexity of distance-sweeping procedures.

Finally, the per-cell nature of the score makes it straightforward to build sample-aware summaries that are robust to imbalanced sample sizes. Large ST datasets often contain samples with widely varying numbers of segmented cells; global, cell-pooled summaries can therefore be dominated by a small number of very large samples. Because our method yields per-cell values that can be aggregated within each sample (e.g. medians, quantiles, or fractions of cells above a high-score threshold), comparisons across experimental groups can be performed at the sample level, mitigating the influence of extreme cell counts and enabling statistical testing that aligns with the experimental unit. Together, these properties, single-cell resolution, multivariate niche scoring, transparent interpretation on standard neighbor graphs, and sample-robust summarization, make the proposed co-localization score a useful addition to the ST analysis toolkit for both exploratory mapping and cohort-level inference.

## 5 Data Availability

### Mouse ovary

Please contact the authors for access the the Mouse ovary dataset[15].

